# Evidence of neutralizing antibodies against SARS-CoV-2 in domestic cats living with owners with a history of COVID-19 in Lima – Peru

**DOI:** 10.1101/2021.05.26.445880

**Authors:** Luis M. Jara, Cusi Ferradas, Francesca Schiaffino, Camila Sánchez-Carrión, Ana Martinez, Alexandra Ulloa, Gisela Isasi-Rivas, Angela Montalván, Luis Guevara Sarmiento, Manolo Fernández, Mirko Zimic

**Affiliations:** Facultad de Medicina Veterinaria y Zootecnia, Universidad Peruana Cayetano Heredia, Lima, Peru; Unidad de Investigación en Enfermedades Emergentes y Cambio Climático (Emerge), Facultad de Salud Pública y Administración, Universidad Peruana Cayetano Heredia, Lima, Peru; Clinica Veterinaria Gatuario, Lima, Peru; Clinica Veterinaria Los Dominicos, Lima, Peru; Farmacológicos Veterinarios (FARVET), Chincha, Peru; Laboratorio de Bioinformática, Biología Molecular y Desarrollos Tecnológicos, Universidad Peruana Cayetano Heredia, Lima, Peru

**Keywords:** COVID-19, SARS-CoV-2, cats, serology, neutralizing antibodies, One Health

## Abstract

SARS-CoV-2 can infect a variety of wild and domestic animals worldwide. Of these, domestic cats are highly susceptible species and potential viral reservoirs. As such, it is important to investigate disease exposure in areas with active community transmission and high disease prevalence. In this report we demonstrate the presence of serum neutralizing antibodies against the receptor binding-domain (RBD) of the SARS-CoV-2 in cats whose owners had been infected with SARS-CoV-2 in Lima, Peru, using a commercial competitive ELISA SARS-CoV-2 Surrogate Virus Neutralization Test. Out of 41 samples, 17.1% (7/41) and 31.7% (13/41) were positive, using the cut-off inhibition value of 30% and 20%, respectively. Not all cats living in a single house had detectable neutralizing antibodies showing that heterogenous exposure and immune among cohabiting animals. This is the first report of SARS-COV-2 exposure of domestic cats in Lima, Peru. Further studies are required to ascertain the prevalence of SARS-COV-2 exposure among domestic cats of Lima, Peru.

## Introduction

The new human coronavirus, SARS-CoV-2, has been shown to mainly infect humans. However, SARS-CoV-2 infection has also been detected in a variety of animals, including wild cats, minks, ferrets, domestic dogs and cats [1-6]. Cats and minks may be considered the most susceptible species because of the higher similarity of the angiotensin-converting enzyme 2 (ACE2) between these species and humans [7]. Although the majority of infected cats are asymptomatic, some animals may develop clinical disease, and the virus can be experimentally transmitted between individuals [8]. Therefore, SARS-CoV-2 could have a direct impact on animal health, while the possibility of cats becoming zoonotic reservoirs has not been totally discarded.

Serological testing is a valuable tool for screening antibody levels associated with pathogen exposure. As with other viral infections, host neutralizing serum antibodies may block the binding of viral proteins to cell surface receptors. In humans, SARS-CoV-2 neutralizing antibodies have been determined to inversely correlate with disease severity and can predict the probability of re-infections [9]. In animals, reported prevalence of neutralizing serum antibodies against SARS-CoV-2 in cats varies, with as low as 0.002% in Germany, 0.2% in Brazil, 5.8% in Italy, and 10.8% in Wuhan, China [10-13]. In Peru, one of the most affected countries by the COVID-19 pandemic, no previous studies have been conducted investigating the seroprevalence or prevalence of SARS-CoV-2 among domestic cats. In this report we demonstrate the presence of serum neutralizing antibodies against the receptor binding-domain (RBD) of the SARS-CoV-2 viral spike protein in cats whose owners confirmed previous infection with SARS-CoV-2.

## Materials and Methods

Blood samples of cats were collected between August 2020 and April 2021 from veterinary centers located in Lima, Peru. All cat owners signed an informed consent authorizing the use of the samples for research purposes. Samples were centrifuged at 3500 rpm for 5 minutes and the serum supernatant was transferred to microcentrifuge tubes and was stored at -20^°^C. Samples from cats whose owners confirmed previous COVID-19 disease (clinical signs with positive IgG/IgM rapid test or qRT-PCR) during veterinary anamnesis were conveniently selected. To test the serum samples for the presence of neutralizing antibodies against the RBD of the viral spike protein, a commercial competitive ELISA SARS-CoV-2 Surrogate Virus Neutralization Test (sVNT) was used (Genscript, New Jersey, USA) according to the manufacturer’s instructions. Percent serum neutralization was calculated as follows: = (1 - OD value of sample / OD value of negative control) × 100%. A cut-off value of 20% and an updated 30% of inhibition were used to establish positivity. The study was approved by the Universidad Peruana Cayetano Heredia Animal Care and Use Ethical Committee (N° 027-08-20).

## Results and Discussion

A total of 41 samples from a serum bank of 700 were selected for screening of serum neutralizing antibodies. The median age of the animals was 12 months (IQR: 8 months – 46 months), 53.7% were female (22/41), and 87.8 % (36/41) were classified as domestic shorthair. 53.7% (22/41) came from the district of Comas while the remainder 46.3% came from Miraflores (5/41), Surco (5/41), San Juan de Miraflores (3/41), Independencia (2/41), San Juan de Lurigancho (1/41), San Luis (1/41), and San Martín de Porres (1/41). Out of the 41 cat samples, 22 cats (53.7%) lived in a single household (household C) in which cats were sampled on two different dates, and 2 cats lived in household D. Age, sex, breed, and district of all animals are shown in **Supplementary Table 1**. Out of the 41 samples, 17.1% (7/41) and 31.7% (13/41) were positive for the presence of serum neutralizing antibodies, using the cut-off value of 30% and 20%, respectively (**Figure 1**). Out of 13 positive samples, 38.4% (5/13) showed clinical signs including sneezing and dyspnea, cough, vomit, or depression. Interestingly, one of the animals with the highest percent neutralization (73.06%) showed all the symptoms described. Only 8 out 22 cats in household C had evidence of serum neutralizing antibodies. This suggests that infection may not be homogenous among cohabiting animals, and this could be associated with other factors such as health state, immunity, proximity to the infected owner(s), among others.

**Figure 1.**
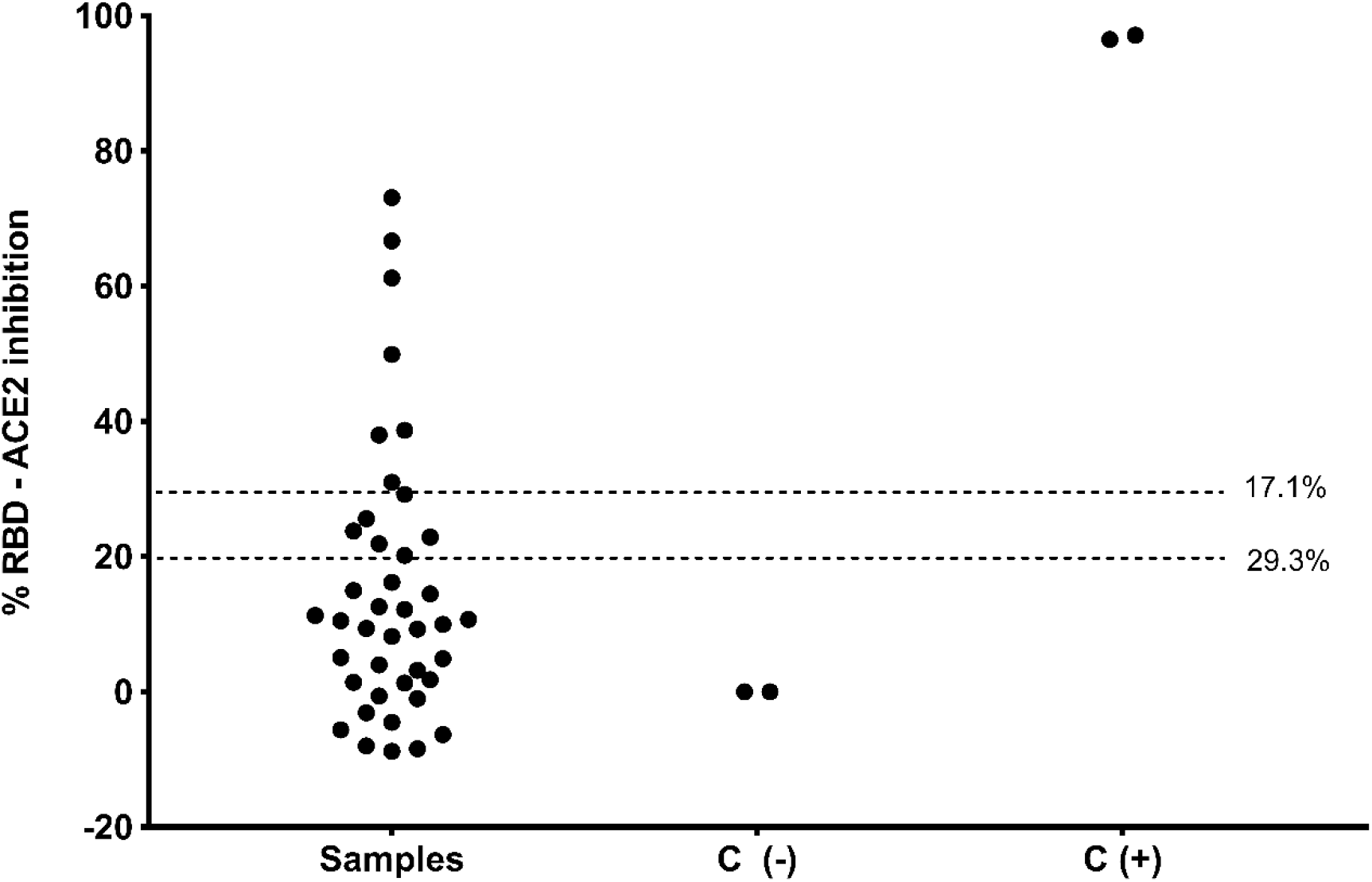
Percent inhibition against SARS-COV-2 receptor binding-domain (RBD) in serum of domestic cats whose owners had a history of COVID-19 (n = 41). C: controls, (-): negative, (+): positive. The 17.1% and 29.3% shows the frequency of cats with neutralizing antibodies with 30% and 20% of cut-off values of inhibition, respectively.

Our results show compelling evidence of SARS-CoV-2 exposure in domestic cats and it is the first report of such an event in Peru. Percent seropositivity in this population of cats is high compared to other studies published, such as that of Italy and Wuhan, China, in which 5.8% of 191 cats 10.8% of 102 cats had neutralizing antibodies, respectively [12, 13]. However, these studies were not exclusively done on a pet population living with COVID-19 infected owners. Ina longitudinal cohort study of pets living with COVID-19 owners, 43.8% of 16 cats developed neutralizing antibodies against SARS-CoV-2 [14]. Limited sample size and a convenience sample do not permit prevalence estimation.

Serum neutralization activity is commonly tested using plaque reduction neutralization tests. However, the commercial assay utilized in this preliminary study has shown a high correlation with serum neutralization activity using plaque reduction and has shown robust internal validity parameters for both humans, cats, dogs and hamster sera [15, 16]. Additionally, this commercial assay offers logistical and biosafety advantages for researchers working in resource-limited settings that do not have access to a BSL-3 containment required for SARS-CoV-2 manipulation.

These animals sought routine veterinary care that was not associated with symptomatic respiratory disease in most of the cases, demonstrating potential asymptomatic infection in cats, and consequently, potential viral reservoirs. In one study, over 25% of households sampled had pets with neutralizing antibodies. Few case studies of natural infection in cats document severe clinical outcomes, and those that have revealed that co-morbidities likely played a contributing factor in illness or death [14].

## Conclusions

It is crucial to monitor SARS-CoV-2 exposure and infection in domestic animals using rapid and affordable point-of-care serological and molecular assays that can be used by veterinarians serving low-income communities. Cats have the potential to serve as sentinels for undetected community transmission, and in this scenario, veterinarians play a key role as first-line responders.

## Ethics Statement

The authors confirm that the ethical policies of the journal, as noted on the journal’s author guidelines page, have been adhered to and the appropriate ethical review committee approval has been received.

## Acknowledgements

Partial funding was received by D43TW007393 from the Emerging Diseases and Climate Change Research Unit (Emerge), Universidad Peruana Cayetano Heredia, Lima, Peru. We would like to acknowledge Dr. Macarena Llalla for helping with the collection of cat serum samples, as well as all other Veterinary supportive staff.

