## Supplementary Table 1 for "Evidence of neutralizing antibodies against SARS-CoV-2 in domestic cats living with owners with a history of COVID-19 in Lima – Peru"

**Supplementary Table 1. Data of cats with serum neutralizing antibodies against SARS-COV-2**

| Household | Sample Date (MM/DD/YY) | Age (months) | Sex | Breed | District | Percent Inhibition (%) |
|---|---|---|---|---|---|---|
| A | 1/20/2021 | 7 | Male | Domestic Shorthair | San Juan de Miraflores | **73.06** |
| A | 1/21/2021 | 28 | Female | Domestic Shorthair | San Juan de Miraflores | **66.69** |
| B | 4/01/2021 | 8 | Female | Domestic Shorthair | San Juan de Miraflores | 3.2 |
| C.1 | 10/9/2020 | 8 | Female | Domestic Shorthair | Comas | **29.15** |
| C.1 | 10/9/2020 | 12 | Female | Domestic Shorthair | Comas | **25.59** |
| C.1 | 10/9/2020 | 11 | Female | Domestic Shorthair | Comas | **23.81** |
| C.1 | 10/09/2020 | 36 | Male | Domestic Shorthair | Comas | **22.88** |
| C.1 | 10/9/2020 | 10 | Female | Domestic Shorthair | Comas | **20.24** |
| C.1 | 10/9/2020 | 5 | Male | Domestic Shorthair | Comas | 10.69 |
| C.1 | 10/9/2020 | 5 | Male | Domestic Shorthair | Comas | 4.92 |
| C.1 | 10/9/2020 | 24 | Male | Domestic Shorthair | Comas | 9.27 |
| C.1 | 10/9/2020 | 12 | Female | Domestic Shorthair | Comas | -3.14 |
| C.1 | 10/9/2020 | 12 | Female | Domestic Shorthair | Comas | -0.64 |
| C.1 | 10/9/2020 | 24 | Female | Domestic Shorthair | Comas | 14.47 |
| C.1 | 10/9/2020 | 10 | Female | Domestic Shorthair | Comas | 16.18 |
| C.1 | 10/9/2020 | 60 | Female | Domestic Shorthair | Comas | 3.99 |
| C.1 | 10/9/2020 | 5 | Female | Domestic Shorthair | Comas | 1.8 |
| C.2 | 10/24/2020 | 9 | Male | Domestic Shorthair | Comas | **61.22** |
| C.2 | 10/24/2020 | 6 | Male | Domestic Shorthair | Comas | **49.87** |
| C.2 | 10/24/2020 | 11 | Male | Domestic Shorthair | Comas | **38.03** |
| C.2 | 10/24/2020 | 8 | Female | Domestic Shorthair | Comas | -8.38 |
| C.2 | 10/24/2020 | 8 | Female | Domestic Shorthair | Comas | -8.75 |
| C.2 | 10/24/2020 | 24 | Female | Domestic Shorthair | Comas | 12.58 |
| C.2 | 10/24/2020 | 7 | Female | Domestic Shorthair | Comas | 8.18 |
| C.2 | 10/24/2020 | 6 | Female | Domestic Shorthair | Comas | -5.59 |
| D | 1/27/2021 | 24 | Female | Domestic Shorthair | Independencia | **38.73** |
| D | 1/27/2021 | 60 | Male | Domestic Shorthair | Independencia | **30.96** |
| F.1 | 8/17/2020 | 84 | Male | Russian Blue | Miraflores | **21.88** |

| | | | | | | |
|---|---|---|---|---|---|---|
| F.2 | 8/31/2020 | 72 | Female | Russian Blue | Miraflores | 9.41 |
| F.2 | 8/31/2020 | 12 | Male | Bengal | Miraflores | 11.26 |
| F.2 | 8/31/2020 | 12 | Male | Bengal | Miraflores | 15.04 |
| G | 2/14/2021 | 46 | Female | Maine Coon | Miraflores | 10.48 |
| H | 12/29/2020 | 96 | Female | Domestic Shorthair | Surco | 5.06 |
| I | 1/13/2021 | 6 | Male | Domestic Shorthair | Surco | -7.98 |
| J | 4/03/2021 | 58 | Male | Domestic Shorthair | Surco | 1.31 |
| K | 1/02/2021 | 64 | Male | Domestic Shorthair | Surco | -4.48 |
| L | 2/06/2021 | 42 | Female | Domestic Shorthair | Surquillo | -6.33 |
| M | 6/02/2021 | 77 | Male | Domestic Shorthair | San Luis | 1.39 |
| N | 1/03/2021 | 124 | Male | Domestic Shorthair | San Juan de Lurigancho | 12.19 |
| O | 10/13/2021 | 63 | Male | Domestic Shorthair | San Martín de Porres | 10.05 |
| P | 2/02/2021 | 9 | Male | Domestic Shorthair | Surco | -1.02 |

Different capital letters represent different households. Household “C” and “F” were sampled in two time points, such that C.1 and F.1 represents the first sampling event and C.2 and F.2 the second one. Percent inhibition (%) in denotes positivity.
